# Extensive membrane lipid remodeling at the host–parasite interface of *Toxoplasma gondii*

**DOI:** 10.64898/2026.11.01.763526

**Authors:** Yuri Nakashima, Sachiko Ishida, Nanaru Mineoka, Tatsunori Masatani, Sayuri Kuriyama, Miho Yoshida, Yoshifumi Nishikawa, Akikazu Fujita

## Abstract

Phosphoinositides regulate diverse membrane trafficking and signaling events in eukaryotic cells, yet their transbilayer distribution remains largely unexplored, particularly at host–parasite membrane interfaces. Here, we determined the membrane leaflet localization of phosphatidylinositol 4-phosphate (PtdIns(4)P) and phosphatidylinositol 4,5-bisphosphate (PtdIns(4,5)P_2_) in *Toxoplasma gondii* using quick-freezing freeze-fracture replica labeling combined with phosphoinositide-specific probes. Both phosphoinositides were predominantly localized to the cytoplasmic leaflet of the parasite plasma membrane, whereas only weak labeling was detected in the inner membrane complex. PtdIns(4)P, but not PtdIns(4,5)P_2_, was additionally detected on intracellular vesicles, while mature micronemes lacked detectable labeling for either phosphoinositide. Surprisingly, both phosphoinositides exhibited nearly symmetric distributions between the two leaflets of the parasitophorous vacuole membrane (PVM), in striking contrast to their asymmetric localization in the parasite plasma membrane. Together with our previous demonstration that phosphatidylserine and phosphatidylethanolamine also lose their transbilayer asymmetry in the PVM, these findings indicate that the PVM undergoes global membrane lipid remodeling that extends from major phospholipids to phosphoinositides. We propose that this extensive lipid reorganization represents a defining structural feature of the apicomplexan PVM and provides a new framework for understanding membrane identity at the host–parasite interface.

## 1. Introduction

The asymmetric distribution of membrane lipids between the two leaflets of biological membranes is a fundamental feature of eukaryotic cells and is essential for the spatial organization and function of cellular membranes [1–3]. In particular, the cytoplasmic leaflet is enriched in phosphatidylserine, phosphatidylethanolamine, and phosphoinositides, which provide a platform for membrane-associated proteins and signaling molecules. Among membrane lipids, phosphoinositides are particularly important because they serve as membrane landmarks that recruit a wide variety of cytosolic effector proteins through specific lipid-binding domains, thereby defining membrane identity and directing intracellular trafficking [4]. Consequently, their asymmetric localization within the cytoplasmic leaflet is crucial for the spatial regulation of membrane-associated signaling events [5].

Among the phosphoinositides, PtdIns(4)P and PtdIns(4,5)P are central regulators of membrane dynamics. In mammalian cells, PtdIns(4)P is enriched primarily at the Golgi apparatus and, to a lesser extent, at the plasma membrane, whereas PtdIns(4,5)P is concentrated almost exclusively in the cytoplasmic leaflet of the plasma membrane, where it regulates endocytosis, exocytosis, actin dynamics, vesicle trafficking, and numerous signaling pathways [4, 6, 7]. Because phosphoinositides function by recruiting cytosolic proteins, their confinement to the cytoplasmic leaflet is generally regarded as a fundamental principle underlying membrane identity in eukaryotic cells.

*Toxoplasma gondii* is an obligate intracellular apicomplexan parasite with a remarkably broad host range, infecting humans and a wide variety of warm-blooded animals [8]. Following invasion, the parasite resides within a parasitophorous vacuole (PV), which is enclosed by the parasitophorous vacuole membrane (PVM) [9, 10]. The PVM constitutes the principal host–parasite interface and plays essential roles in nutrient acquisition, host immune evasion, intracellular survival, and parasite replication [11–13]. Although numerous PVM-associated proteins have been identified, remarkably little is known about the molecular organization of the membrane lipids that constitute this unique host–parasite interface.

Recent studies have established important roles for phosphoinositide metabolism in apicomplexan parasites, including *T. gondii* and *Plasmodium falciparum* [14]. In *P. falciparum*, phosphatidylinositol 4-kinase (PI4K) has been identified as an essential and druggable target, and pharmacological inhibition of PI4K disrupts parasite development and replication [15, 16]. In *T. gondii*, phosphoinositide-dependent pathways have likewise been implicated in essential cellular processes, including membrane trafficking and organelle biogenesis [17, 18]. In *T. gondii*, PtdIns(4)P has been implicated in Golgi and post-Golgi membrane trafficking and in the biogenesis and secretion of specialized secretory organelles. Recent studies have demonstrated PtdIns(4)P pools in the Golgi/trans-Golgi network and post-Golgi compartments and shown that disruption of PtdIns(4)P signaling impairs secretory organelle biogenesis and exocytosis [19]. PtdIns(4,5)P has also been implicated in processes essential for parasite motility and host-cell invasion, which depend on coordinated regulation of the parasite plasma membrane and actin–myosin cytoskeleton. Moreover, PI-PLC hydrolyzes PtdIns(4,5)P to generate the second messengers IP and diacylglycerol (DAG), thereby stimulating intracellular Ca² release [20, 21]. Consistent with this pathway, ethanol-induced microneme secretion is accompanied by intracellular Ca² elevation, likely mediated through PI-PLC-derived IP [22–24]. Despite these important biological functions, to date, the subcellular localization of phosphoinositides in *T. gondii* has been investigated predominantly by fluorescence microscopy using genetically encoded phosphoinositide-binding probes [14, 25]. However, these approaches do not provide information on the transbilayer distribution of phosphoinositides between the cytoplasmic and luminal/exoplasmic leaflets of individual membranes.

Although fluorescent lipid-binding probes have greatly advanced our understanding of phosphoinositide biology, they possess important limitations. Overexpression of phosphoinositide-binding domains can compete with endogenous lipid effectors and perturb normal phosphoinositide-dependent processes [7, 26]. For example, overexpression of the

PtdIns(3)P-binding Fab1, YOTB, Vac1, and EEA1 (FYVE) or Phox homology (PX) domains interferes with phagosome maturation [14, 27]. Furthermore, fluorescence microscopy visualizes phosphoinositides only on the cytoplasmic surface and cannot distinguish between the two membrane leaflets. Conventional electron microscopy also lacks sufficient membrane contrast and spatial resolution to assign lipids to individual leaflets.

Quick-freezing and freeze-fracture replica labeling (QF-FRL) overcomes these limitations by physically immobilizing membrane lipids and allowing direct visualization of their distribution on individual membrane leaflets with nanometer resolution [28–31]. Using this approach, we previously demonstrated that the PVMs of both *T. gondii* and *P. falciparum* lose the characteristic transbilayer asymmetry of phosphatidylserine and phosphatidylethanolamine, exhibiting nearly symmetric lipid distributions between the two membrane leaflets [32]. These findings suggested that the PVM does not simply inherit the membrane organization of the host plasma membrane but instead undergoes extensive membrane lipid remodeling during or after parasite invasion.

Whether this remarkable remodeling also extends to phosphoinositides has remained completely unknown. Because phosphoinositides function as membrane landmarks that recruit numerous cytosolic effector proteins, loss of their transbilayer asymmetry would fundamentally alter membrane identity and could profoundly influence host–parasite interactions.

In the present study, we applied QF-FRL together with phosphoinositide-specific probes to determine, for the first time, the membrane leaflet distribution of PtdIns(4)P and PtdIns(4,5)P in *T. gondii*. We found that both phosphoinositides are predominantly localized to the cytoplasmic leaflet of the parasite plasma membrane but become symmetrically distributed between the two leaflets of the PVM. Unexpectedly, PtdIns(4)P was considerably more abundant than PtdIns(4,5)P in the parasite plasma membrane, opposite to the phosphoinositide composition characteristic of mammalian cells. Together with our previous findings on phosphatidylserine and phosphatidylethanolamine [32], these results demonstrate that membrane lipid remodeling in the PVM extends beyond major phospholipids to phosphoinositides. We therefore propose that global membrane lipid remodeling represents a defining structural feature of the apicomplexan parasitophorous vacuole membrane and constitutes a previously unrecognized principle governing membrane organization at the host–parasite interface.

## 2. Materials and Methods

### 2.1. Probes

Complementary DNA (cDNA) generated by RT-PCR of HEK293 cell total RNA was used as a template to clone the PH domains of OSBP. Recombinant GST fusion proteins containing the OSBP PH domain (GST-OSBP-PH) and phospholipase C (PLC)-δ1 PH domain (GST-PLCδ1-PH) were expressed in *Escherichia coli* and purified as previously described (Fig. S1 for GST-OSBP-PH) [29] [33, 34]. The PH domain of PLC-δ1 was obtained from the GST-PLCδ1-PH domain fusion protein via cleavage with PreScission protease (GE Healthcare Life Sciences, Pittsburgh, PA, USA) according to the manufacturer’s protocol. The secondary rabbit anti-GST antibody was purchased from Bethyl Laboratories (Montgomery, TX, USA). 10 nm gold particle-conjugated anti-rabbit IgG antibody were purchased from Jackson ImmunoResearch (West Grove, PA, USA).

### 2.2. Human foreskin fibroblast-1

Human foreskin fibroblast-1 (HFF-1) was cultured in Dulbecco’s modified Eagle’s medium (DMEM; GIBCO, Grand Island, NE, USA) supplemented with 10% fetal calf serum, 50 U/mL penicillin, and 0.05 mg/mL streptomycin at 37 °C under 5% CO_2_.

### 2.3. Parasite lines and culture

*T. gondii* tachyzoites of the PLK strain (kindly provided by Dr. Kami Kim, Albert Einstein College of Medicine, NY, USA) were maintained in HFF-1 cultured in Dulbecco’s modified Eagle’s medium (DMEM; GIBCO, Grand Island, NE, USA) supplemented with 10% fetal calf serum, 50 U/mL penicillin, and 0.05 mg/mL streptomycin at 37 °C under 5% CO_2_. For experiments, *T. gondii* tachyzoites were inoculated into confluent HFF-1 monolayers and incubated for 2–3 days post-infection before sample preparation. *T. gondii* tachyzoites were prepared by washing infected cells with cold phosphate-buffered saline (PBS) and detaching them with a cell scraper. Cell pellets were re-suspended in DMEM and passed through a 27-gauge needle five times and then through a 5.0 μm-pore filter (Millipore, Bedford, MA, USA). Purified tachyzoites were centrifuged at 5,000 ×*g* for 5 min to produce a pellet of cells and used for quick-freezing immediately.

### 2.4. Quick-freezing and freeze-fracture

For metal-sandwich freezing of *T. gondii*, purified tachyzoites were collected by centrifugation, and a small volume of the resulting pellet was placed on a copper foil, covered with a thin gold foil (∼4 mm² in area; 20 μm thick), and rapidly frozen by pressing it between two gold-plated copper blocks precooled in liquid nitrogen [33, 35]. Purified tachyzoites were used to examine the lipid distribution in the parasite plasma membrane and intracellular organelle membranes.

For high-pressure freezing, *T. gondii*-infected HFF-1 cells cultured on gold foil (∼4 mm² in area; 20 μm thick) were sandwiched between flat aluminum discs (Engineering Office M. Wohlwend, Sennwald, Switzerland) and frozen according to the manufacturer’s instructions [36]. These samples were used to examine the lipid distribution in the parasitophorous vacuole membrane (PVM) without isolating the parasites from the host cells.

For metal-sandwich freezing of HFF-1 cells, cells cultured on a small gold foil (∼4 mm² in area; 20 μm thick) were inverted onto prewarmed 10% gelatin on a copper foil, with the cell side facing down, and rapidly frozen using the metal-sandwich method described above [33, 35]. These samples were used to examine the lipid distribution in the HFF-1 cell plasma membrane.

The frozen specimens were transferred to the cold stage of a Balzers BAF400 apparatus (Bal-Tec AG, Lichtenstein) and fractured at −130 °C under a vacuum of ∼1 × 10^−6^ ^m^bar. Replicas were produced by electron-beam evaporation in three steps: carbon (C; ∼2 nm thickness) at an angle of 90°, platinum–carbon (Pt/C; 1–2 nm thickness) at an angle of 45°, and C (10–20 nm thickness) at an angle of 90° to the specimen surface [30]. The deposition thickness was adjusted using a crystal-thickness monitor (EM QSG100, Leica).

The thawed specimens were treated with 2.5% sodium dodecyl sulfate (SDS) in 0.1 M Tris-HCl (pH 8.0) overnight at 60–70 °C, and the replicas were stored in 50% buffered glycerol at −30 °C until used.

### 2.5. Labeling and EM observation

Probe labeling was performed as previously described [30]. Briefly, after rinsing, the freeze-fracture replicas were blocked with PBS containing 2% cold-water fish-skin gelatin at room temperature for 30 min. The replicas were then incubated overnight at 4 °C with the GST-OSBP-PH (40 ng/mL), or GST-PLCδ1-PH (32 ng/mL) diluted in PBS containing 2% cold-water fish-skin gelatin. To block the non-specific binding of GST-OSBP-PH to PtdIns(4,5)P_2_, the replicas were pretreated with the PH domain (1 mg/mL) of PLC-δ1 in the blocking solution [34, 37]. After four washes with PBS containing 0.1% bovine serum albumin (BSA), the replicas were incubated at 37 °C for 30 min with rabbit anti-GST antibody (5 μg/mL), followed by incubation with 50-fold diluted 10 nm gold-conjugated anti-anti-rabbit in PBS containing 2% cold-water fish-skin gelatin. The replicas were transferred to Formvar-coated grids and examined using an H7650 electron microscope (HITACHI, Tokyo, Japan) operated at 80 kV.

### 2.6. Statistical analysis

EM and fluorescence images obtained from at least three independent experiments were used for the analyses. The number of colloidal gold particles was counted manually, and the areas were measured using ImageJ software [38]. The labeling density in the selected structure was calculated by dividing the number of colloidal gold particles by the area. For each structure, the labeling density was measured in more than ten different randomly captured micrographs. Statistical differences between the samples were analyzed using Student’s *t*-test.

## 3. Results

### 3.1. Validation of GST-OSBP-PH labeling for PtdIns(4)P using artificial liposomes

To examine the relationship between PtdIns(4)P content and GST–OSBP-PH labeling, we analyzed artificial liposomes containing increasing amounts of PtdIns(4)P by QF-FRL. Our previous study demonstrated that GST–PLCδ1-PH preferentially labels PtdIns(4,5)P_2_-containing liposomes under the conditions examined [29]. In the present study, GST–OSBP-PH labeling density increased with increasing PtdIns(4)P content (Fig. S1). The relationship was approximately linear over the lower concentration range examined, although a tendency toward saturation was observed at the highest concentration tested.

These results demonstrate that GST–OSBP-PH labeling responds to changes in PtdIns(4)P content in artificial liposomes under the conditions used. However, the labeling efficiencies of GST–OSBP-PH and GST–PLCδ1-PH have not been calibrated against each other under equivalent conditions. Moreover, the lipid composition and biophysical properties of cellular membranes differ from those of artificial liposomes. Therefore, these experiments do not establish a direct quantitative relationship between the labeling densities obtained with the two probes or permit direct comparison of the relative abundance of PtdIns(4)P and PtdIns(4,5)P in cellular membranes.

### 3.2. Localization of PtdIns(4)P in Toxoplasma gondii

To determine the subcellular distribution of phosphatidylinositol 4-phosphate (PtdIns(4)P), freeze-fracture replicas of *T. gondii* tachyzoites were labeled with GST–OSBP-PH, a specific probe for PtdIns(4)P. Dense labeling was predominantly observed on the P-face (PF; cytoplasmic leaflet) of the parasite plasma membrane (Fig. 1). PtdIns(4)P labeling was also detected on the membranes of a subset of intracellular vesicles (Fig. 2A), although some vesicles displayed substantially lower labeling densities (arrowheads in Fig. 2A, Table 1). In contrast, small apical vesicles immediately beneath the plasma membrane exhibited little or no detectable labeling (arrows in Fig. 2B, Table 1). Based on their size, morphology, and localization, these vesicles are consistent with micronemes [39]. Weak but reproducible labeling was also observed on the E-face (EF) of the inner membrane complex (IMC), although the labeling density was markedly lower than that of the plasma membrane (Figs. 1 and 7). Quantitative analysis confirmed that PtdIns(4)P was highly enriched on the PF of the plasma membrane, whereas only low levels were detected on the IMC (Fig. 7).

**Fig. 1.**
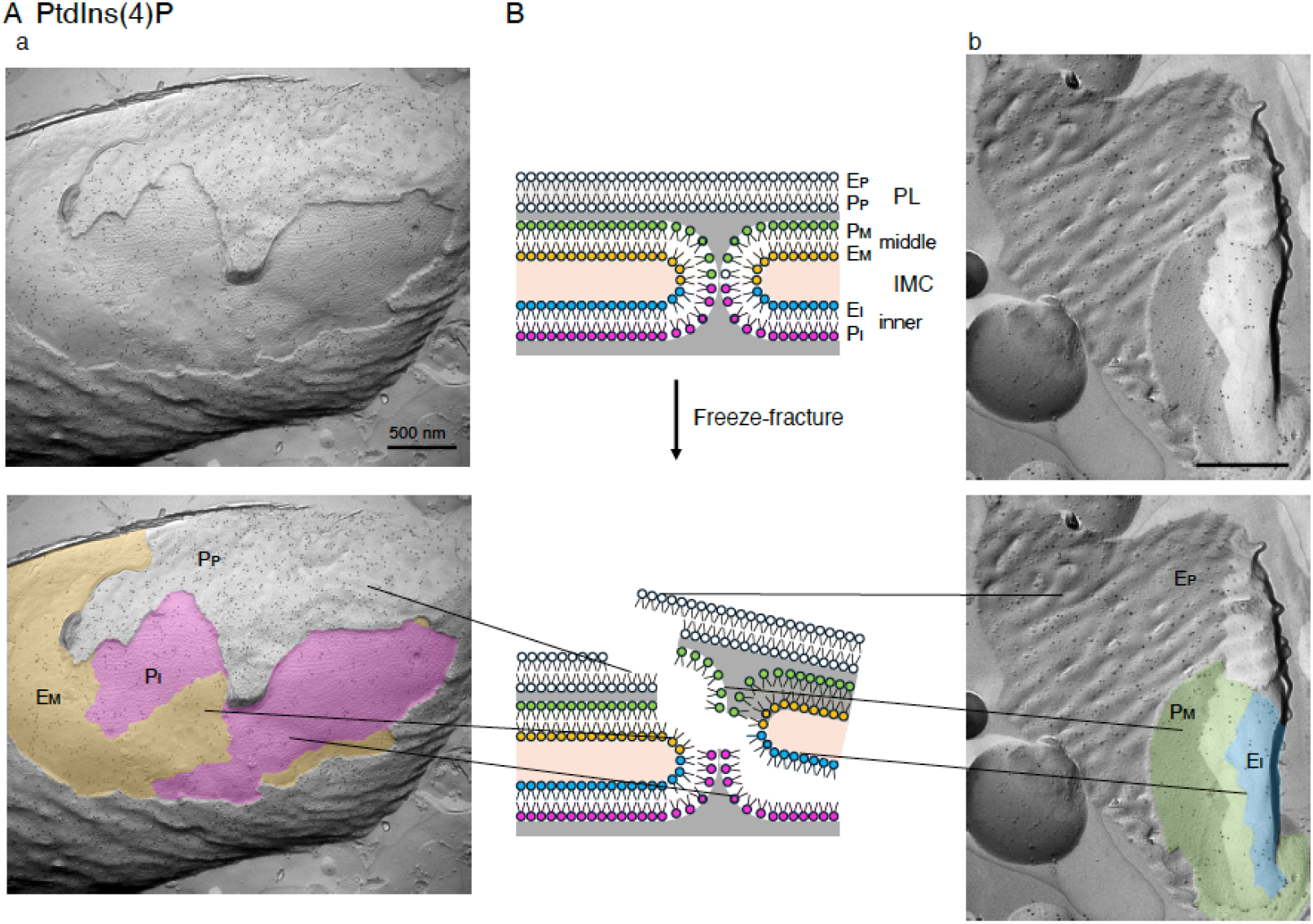
Enrichment of PtdIns(4)P in the cytoplasmic leaflet of the plasma membrane in *T. gondii* tachyzoites. Replicas of *T. gondii* tachyzoites were labeled with a GST–OSBP-PH fusion probe that specifically recognizes PtdIns(4)P. (A) Numerous PtdIns(4)P signals were predominantly detected on the protoplasmic fracture (PF) of the plasma membrane, whereas only sparse labeling was observed on the exoplasmic fracture (EF). PtdIns(4)P labeling was also minimal on both the middle inner membrane complex (IMC; E_M_ and P_M_, yellow and green) and the inner IMC (E_I_ and P_I_, blue and pink) membranes. Scale bar: 500 nm. (B) Schematic representation of the freeze-fracture method and the corresponding fracture planes in *T. gondii* tachyzoites. PL, plasma membrane; IMC, inner membrane complex.

**Fig. 2.**
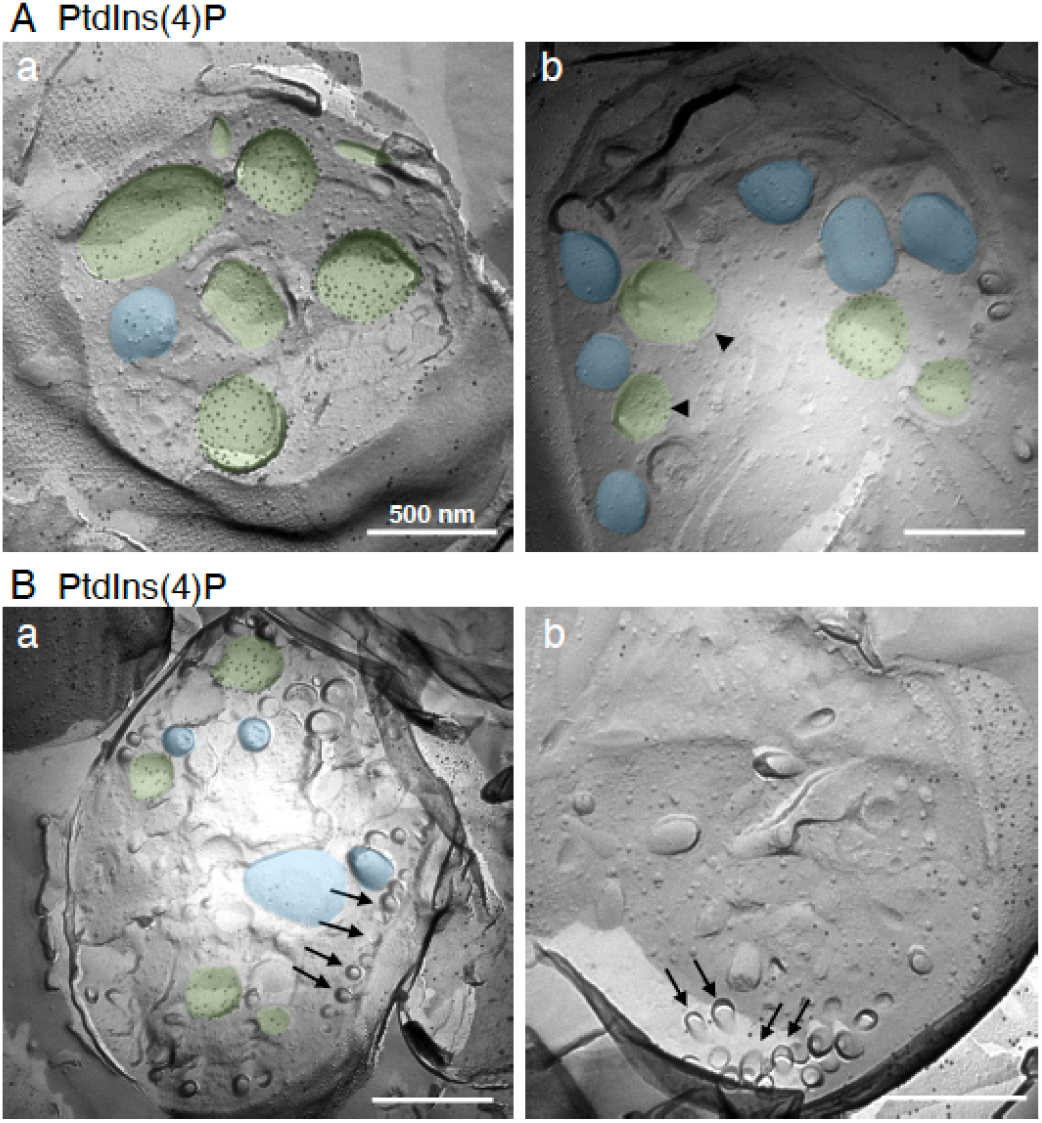
Distribution of PtdIns(4)P in the membranes of the intracellular vesicles. PtdIns(4)P labeling was predominantly detected on the PF (green), but not on the EF (blue), of intracellular vesicular membranes, although not all vesicles exhibited abundant PtdIns(4)P signals (arrowheads in panel Aa). No PtdIns(4)P labeling was observed on the membranes of micronemes located at the periphery of *T. gondii* tachyzoites (arrows in panel B). Scale bar: 500 nm.

**Table 1.** Average sizes of PtdIns(4)P-positive and -negative intracellular vesicles in *T. gondii* tachyzoites. The mean diameter of the small intracellular vesicles lacking PtdIns(4)P labeling was significantly smaller than that of the PtdIns(4)P-positive vesicles in *T. gondii* tachyzoites. *t*-test: \*\*\**p* < 0.001; n.s., not significant.

### 3.3. Membrane leaflet distribution of PtdIns(4)P in the parasitophorous vacuole membrane

We next examined the membrane leaflet distribution of PtdIns(4)P in the PVM of *T. gondii*-infected HFF-1 cells (Fig. 3). Unlike the parasite plasma membrane, PtdIns(4)P labeling was detected on both the PF and EF of the PVM. Quantitative analysis revealed no significant difference in labeling density between the two membrane leaflets, indicating that PtdIns(4)P is distributed nearly symmetrically across the PVM (Fig. 7).

**Fig. 3.**
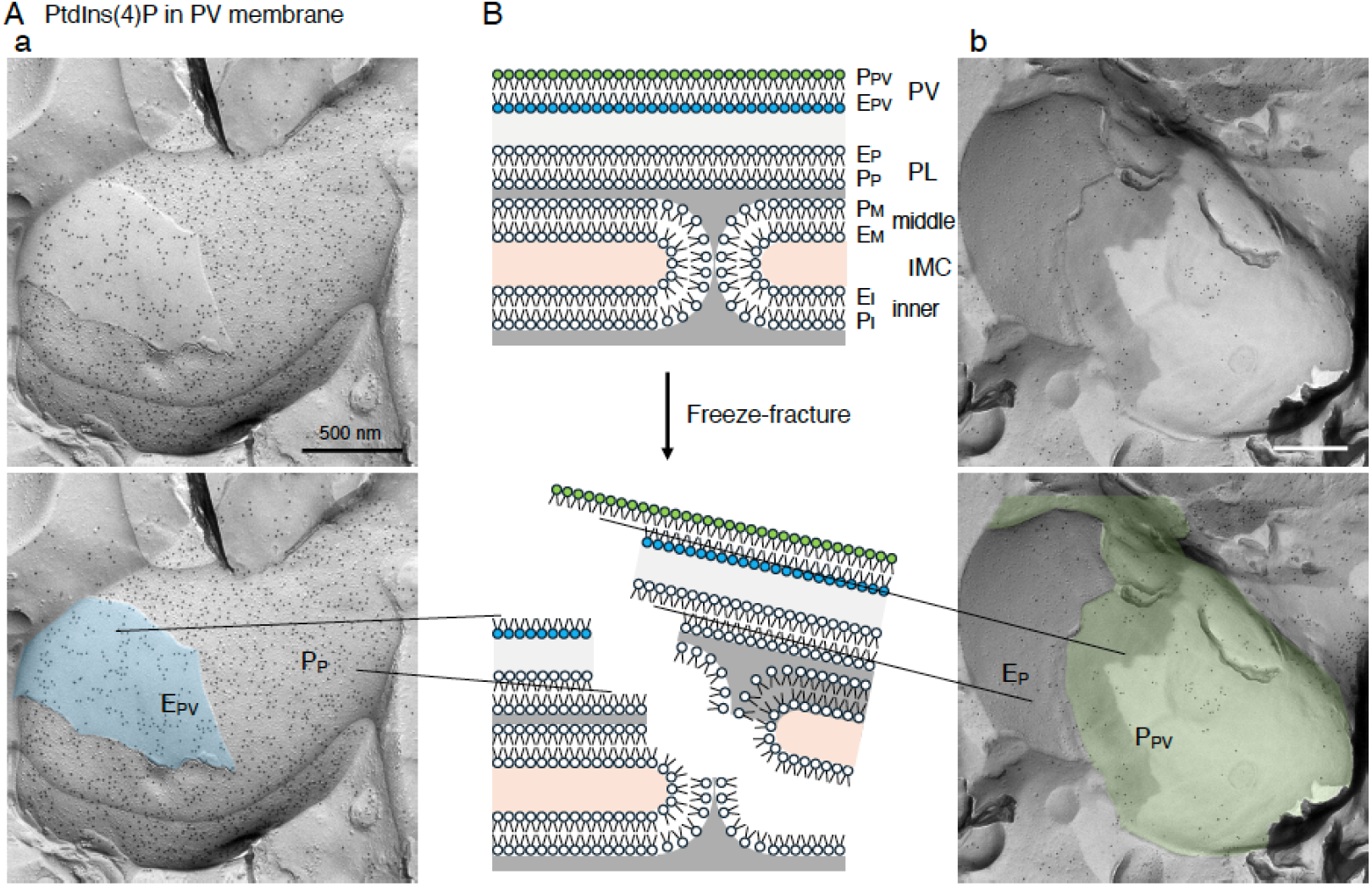
Symmetrical localization of PtdIns(4)P on the EF and PF of the parasitophorous vacuolar membrane. PtdIns(4)P was also localized to the parasitophorous vacuolar (PV) membrane of *T. gondii*. Labeling was symmetrically distributed on both the EF (blue, E_PV_) and PF (green, P_PV_) of the PV membrane.

### 3.4. Localization of PtdIns(4,5)P_2_ in Toxoplasma gondii

The subcellular distribution of phosphatidylinositol 4,5-bisphosphate (PtdIns(4,5)P_2_) was examined using GST–PLCδ1–PH. Similar to PtdIns(4)P, PtdIns(4,5)P_2_ was predominantly localized to the PF of the parasite plasma membrane (Fig. 4). Weak labeling was also detected on the IMC, whereas only sparse labeling was observed on intracellular vesicles (Fig. 5), including micronemes (arrows in Fig. 5). Quantitative analysis confirmed that intracellular vesicles contained substantially lower levels of PtdIns(4,5)P_2_ than PtdIns(4)P (Fig. 7).

**Fig. 4.**
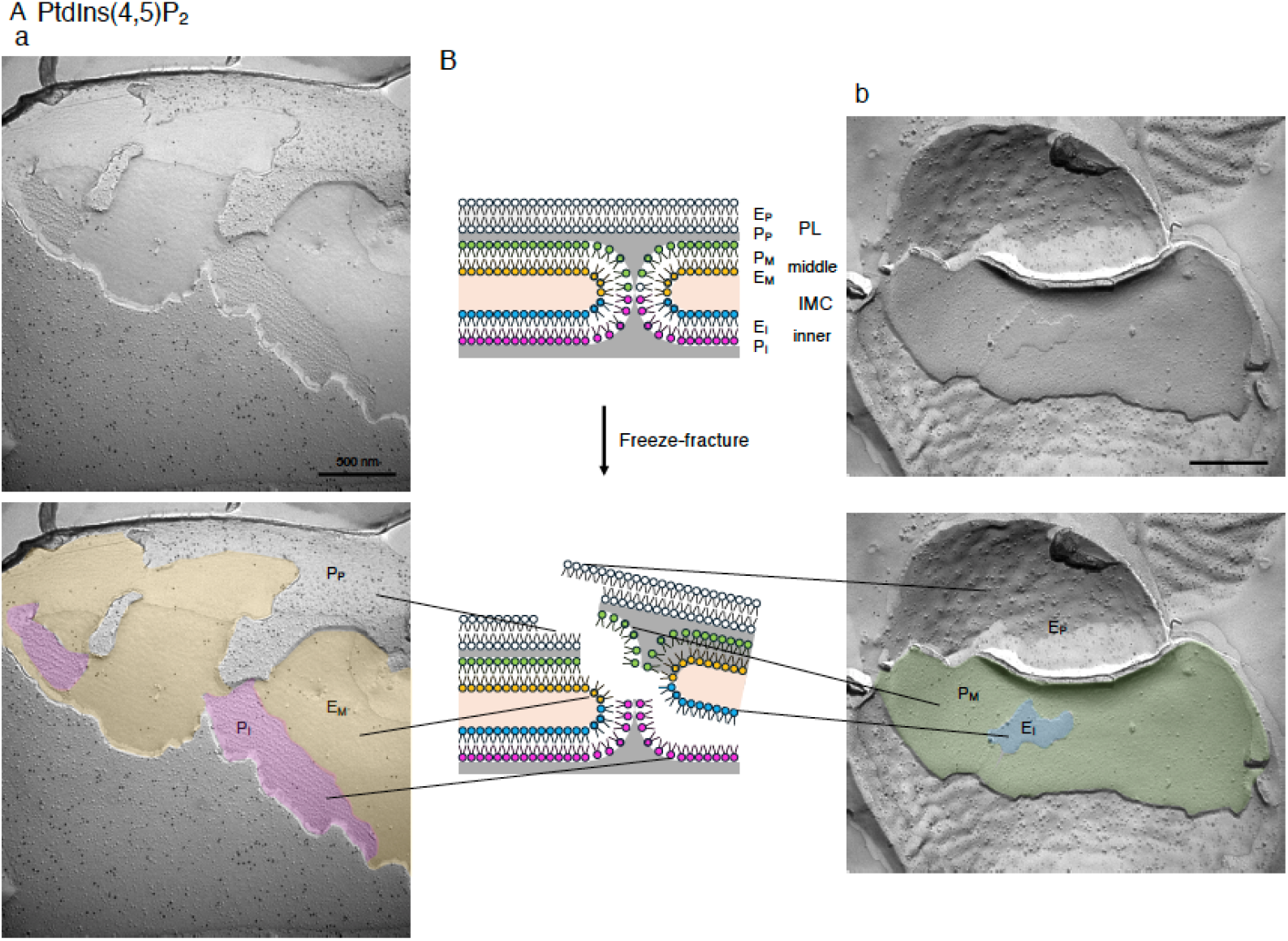
Localization of PtdIns(4,5)P_2_ in the plasma membrane and IMC membranes of *T. gondii*. Replicas of *T. gondii* tachyzoites were labeled with a GST–PLCδ1-PH fusion probe that specifically recognizes PtdIns(4,5)P_2_. (A) A small number of PtdIns(4,5)P_2_ signals were predominantly detected on the PF of the plasma membrane, whereas labeling on the EF was sparse. PtdIns(4,5)P labeling was also minimal on both the middle IMC (E_M_ and P_M_, yellow and green) and the inner IMC (E_I_ and P_I_, blue and pink) membranes. Scale bar: 500 nm. (B) Schematic representation of the freeze-fracture method and the fracture planes in *T. gondii* tachyzoites. PL, plasma membrane; IMC, inner membrane complex.

**Fig. 5.**
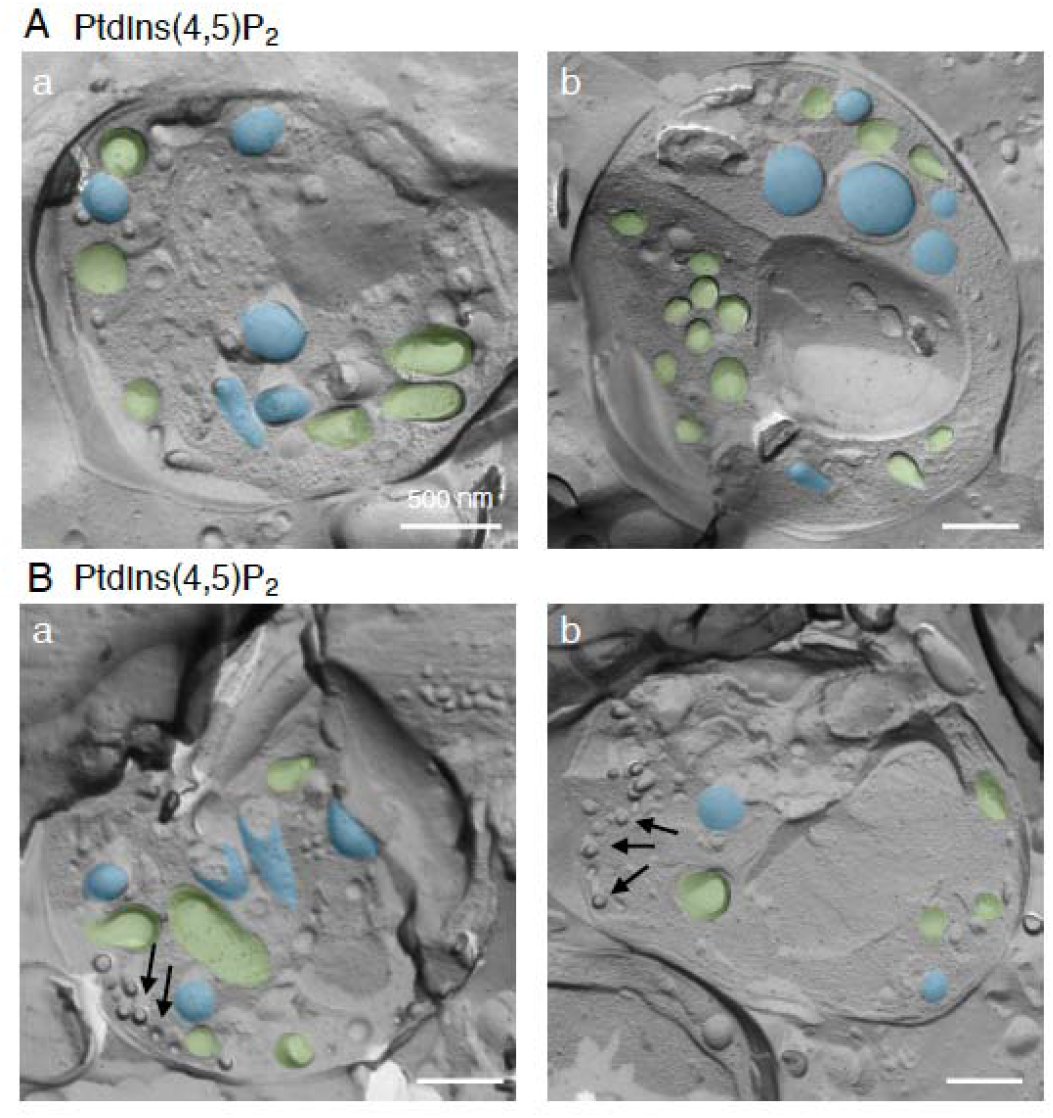
Localization of PtdIns(4,5)P_2_ in the membrane of the intracellular vesicles in *T. gondii*. PtdIns(4,5)P_2_ labeling was predominantly detected on the PF (green), rather than the EF (blue), of intracellular vesicular membranes, although not all vesicles exhibited abundant labeling (arrowheads in panel Aa). Scale bar: 500 nm.

### 3.5. Membrane leaflet distribution of PtdIns(4,5)P_2_ in the parasitophorous vacuole membrane

PtdIns(4,5)P_2_ was also detected in the PVM of *T. gondii*-infected cells (Fig. 6). Although the overall labeling density was considerably lower than that of PtdIns(4)P, labeling was observed on both membrane leaflets. Quantitative analysis demonstrated no significant difference between the PF and EF, indicating that PtdIns(4,5)P_2_ also exhibits a nearly symmetric distribution across the PVM (Fig. 7).

**Fig. 6.**
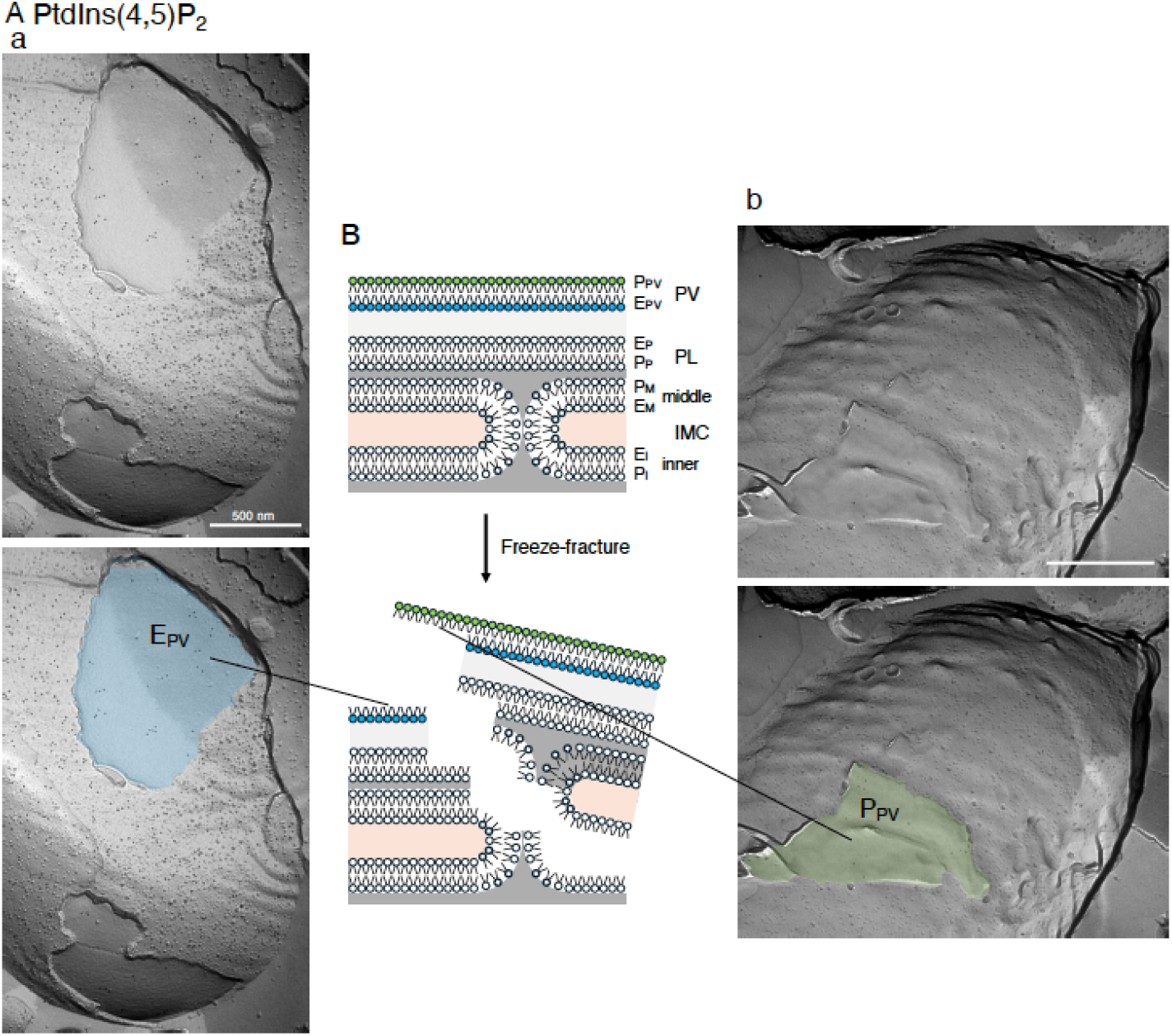
Localization of PtdIns(4,5)P_2_ on both the EF and PF in the parasitophorous vacuolar membrane of *T. gondii*. PtdIns(4,5)P was also localized to the parasitophorous vacuolar (PV) membrane of *T. gondii*. Labeling was symmetrically distributed on both the EF (blue, E_PV_) and PF (green, P_PV_) of the PV membrane.

### 3.6. Comparison of phosphoinositide labeling densities in the parasite and host-cell plasma membranes

Quantitative analysis revealed differences in the labeling densities obtained with GST–OSBP-PH and GST–PLCδ1-PH in the plasma membranes of *T. gondii* and host cells (Fig. 7). In the *T. gondii* plasma membrane, the labeling density obtained with GST–OSBP-PH was higher than that obtained with GST–PLCδ1-PH. In contrast, in the host-cell plasma membrane, the labeling density obtained with GST–PLCδ1-PH was higher than that obtained with GST–OSBP-PH (Fig. 7). Because the two probes have different lipid-binding properties and their labeling efficiencies were not calibrated against each other under equivalent conditions, these differences in labeling density cannot be directly interpreted as differences in the relative abundance of PtdIns(4)P and PtdIns(4,5)P_2_.

**Fig. 7.**
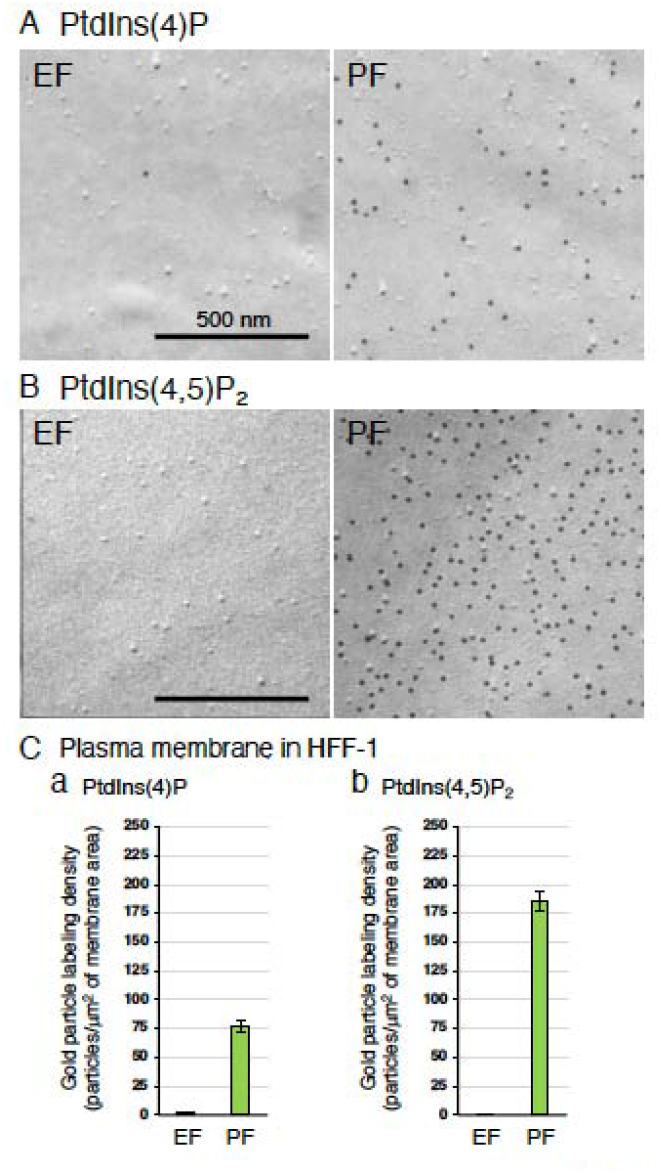
Leaflet-specific distribution and labeling density of PtdIns(4)P and PtdIns(4,5)P_2_ in the plasma membrane of HFF-1 cells. (A) QF-FRL images showing the labeling of PtdIns(4)P on the exoplasmic leaflet (EF) and protoplasmic leaflet (PF) of the HFF-1 cell plasma membrane using GST–OSBP-PH. (B) QF-FRL images showing the labeling of PtdIns(4,5)P_2_ on the EF and PF of the HFF-1 cell plasma membrane using GST–PLCδ1-PH. (C) Quantification of labeling densities for PtdIns(4)P (a) and PtdIns(4,5)P_2_ (b) on the EF and PF of the HFF-1 cell plasma membrane. Labeling density is expressed as the number of colloidal gold particles per μm² of membrane area. Data are presented as the mean ± SEM. The numbers of membrane profiles analyzed were as follows: PtdIns(4)P, EF (n = 18) and PF (n = 24); PtdIns(4,5)P_2_, EF (n = 12) and PF (n = 20). No statistical comparisons were performed.

### 3.7. Asymmetric distribution of phosphoinositides in the host-cell plasma membrane

To validate the specificity of the labeling, phosphoinositide distribution was also examined in the plasma membrane of uninfected HFF-1 cells. Both PtdIns(4)P and PtdIns(4,5)P_2_ were strongly enriched on the PF, whereas only sparse labeling was detected on the EF (Fig. 7), confirming the characteristic transbilayer asymmetry of phosphoinositides in mammalian plasma membranes.

### 3.8. Summary of phosphoinositide distribution

A quantitative summary of phosphoinositide localization is presented in Fig. 8. PtdIns(4)P was highly enriched in the parasite plasma membrane and additionally detected on a subset of intracellular vesicles and the IMC, whereas PtdIns(4,5)P was largely confined to the plasma membrane, with only weak labeling of intracellular membranes. In contrast to the asymmetric distribution observed in both the parasite and host-cell plasma membranes, both phosphoinositides exhibited nearly symmetric distributions across the PVM.

**Fig. 8.**
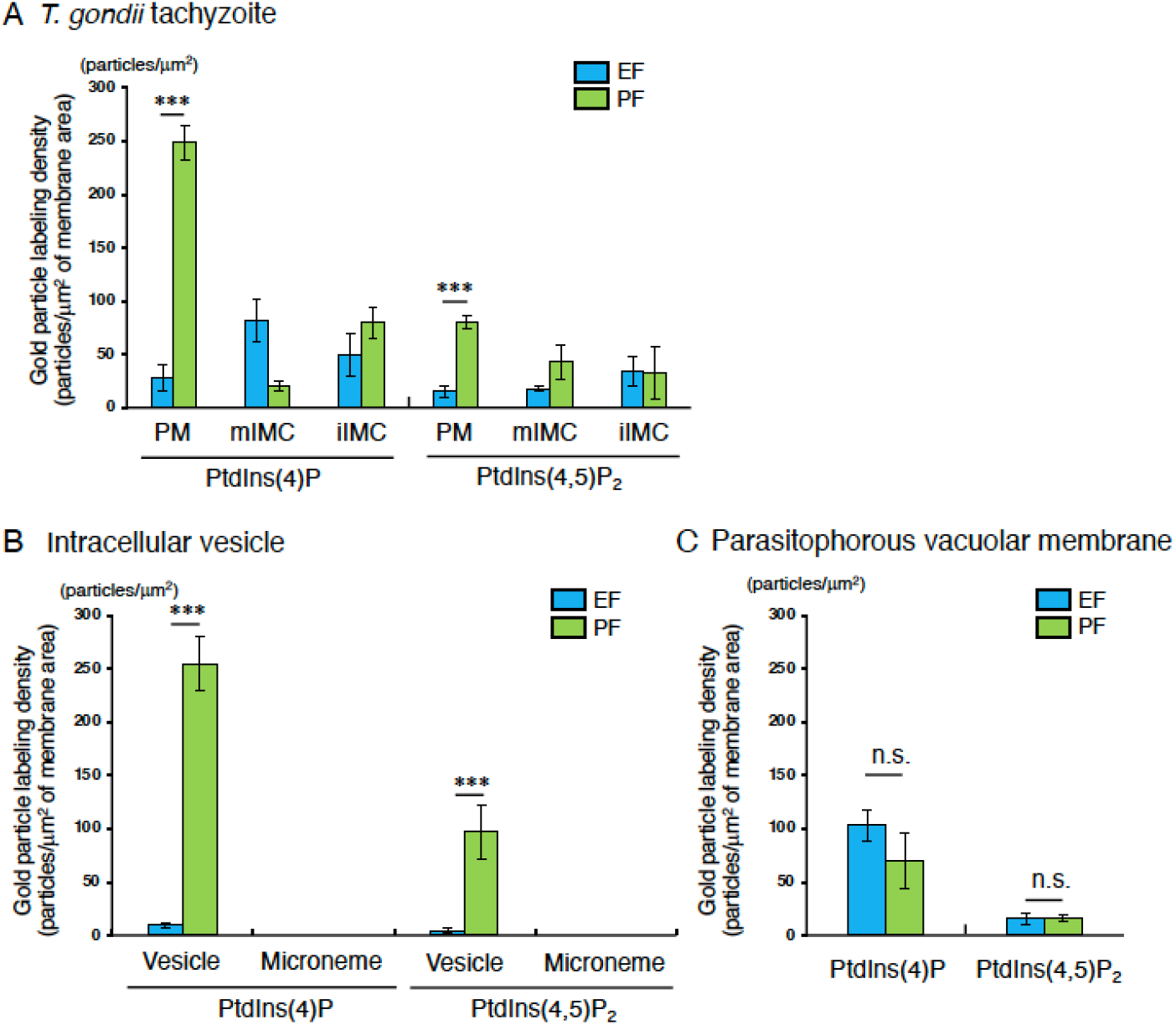
Higher labeling density of PtdIns(4)P than PtdIns(4,5)P_2_ in the PF of *T. gondii* membranes. Labeling densities of PtdIns(4)P and PtdIns(4,5)P_2_ were quantified in the plasma membrane and inner membrane complex (IMC) (A), intracellular vesicles and micronemes (B), and the parasitophorous vacuole (PV) membrane (C) of *T. gondii*. Both phosphoinositides were preferentially detected on the PF relative to the EF of the parasite plasma membrane and intracellular vesicles, whereas similar labeling densities were observed on the EF and PF of the PV membrane. Notably, PtdIns(4)P labeling density was consistently higher than that of PtdIns(4,5)P_2_ in the examined membrane compartments. Labeling density was expressed as the number of colloidal gold particles per μm² of membrane area. PM: plasma membrane, mIMC: middle IMC, iIMC: inner IMC. Data are presented as the mean ± SEM. The numbers of membrane profiles analyzed were as follows: (A) PtdIns(4)P, EF (n = 31) and PF (n = 32) in the PM, EF (n = 35) and PF (n = 32) in the mIMC, EF (n = 37) and PF (n = 31) in the iIMC; PtdIns(4,5)P_2_, EF (n = 34) and PF (n = 37) in the PM, EF (n = 31) and PF (n = 38) in the mIMC, EF (n = 32) and PF (n = 31) in the iIMC. (B) PtdIns(4)P, EF (n = 33) and PF (n = 31) in the vesicles, EF (n = 31) and PF (n = 45) in the micronemes; PtdIns(4,5)P_2_, EF (n = 31) and PF (n = 34) in the vesicles, EF (n = 40) and PF (n = 48) in the micronemes. (C) PtdIns(4)P, EF (n = 38) and PF (n = 37) in the parasitophorous membranes; PtdIns(4,5)P_2_, EF (n = 31) and PF (n = 32) in the parasitophorous membranes. n. s.: not significant, *t*-test, ***, *p* < 0.001

## 4. Discussion

The present study provides a membrane leaflet-specific ultrastructural analysis of PtdIns(4)P and PtdIns(4,5)P_2_ in *Toxoplasma gondii*. Using QF-FRL combined with phosphoinositide-binding probes, we found that both phosphoinositides were predominantly localized to the cytoplasmic leaflet of the parasite plasma membrane, whereas labeling was detected on both leaflets of the PVM (Fig. 8). In addition, PtdIns(4)P labeling was detected on a subset of intracellular vesicles, whereas PtdIns(4,5)P_2_ labeling was sparse on intracellular membranes. These findings reveal distinct subcellular localization patterns of the two phosphoinositides and demonstrate that their transbilayer distributions differ between the parasite plasma membrane and the PVM.

An important consideration in interpreting these results is that gold-particle labeling densities obtained with different lipid-binding probes do not necessarily provide a direct measure of the relative abundance of their target lipids. Our liposome experiments demonstrated a concentration-dependent increase in GST–OSBP-PH labeling with increasing PtdIns(4)P content (Fig. S1B), although the relationship showed a tendency toward saturation at higher concentrations. Furthermore, the labeling efficiencies of GST–OSBP-PH and GST–PLCδ1-PH were not calibrated against each other under equivalent conditions, and the molecular and biophysical environments of cellular membranes differ from those of artificial liposomes. Therefore, differences in labeling densities obtained with these probes in the parasite and host-cell plasma membranes (Figs. 7 and 8A) cannot be directly interpreted as differences in the relative abundance of PtdIns(4)P and PtdIns(4,5)P_2_. Accordingly, our conclusions focus on the subcellular and transbilayer distributions revealed by each probe rather than on a direct quantitative comparison of the two phosphoinositides.

In mammalian cells, PtdIns(4)P is present at the plasma membrane, where it contributes to phosphoinositide metabolism and serves as a precursor for PtdIns(4,5)P_2_ synthesis, although its distribution and functions extend beyond this role [4, 6, 7]. In apicomplexan parasites, phosphoinositide metabolism is essential for parasite biology, with studies implicating PI4KIIIβ in *Plasmodium* [15] and PtdIns(4)P signaling in *T. gondii* [14]. The present findings extend this body of work by defining the leaflet-specific distribution of PtdIns(4)P and PtdIns(4,5)P_2_ at the ultrastructural level in *T. gondii*. In particular, the predominant localization of both phosphoinositides to the cytoplasmic leaflet of the parasite plasma membrane provides a spatial framework for investigating their roles in membrane-associated signaling and lipid metabolism. However, the present data do not establish the relative abundance of the two phosphoinositides or determine their respective contributions to these processes.

The most important finding of the present study is that both PtdIns(4)P and PtdIns(4,5)P_2_ were detected on both leaflets of the PVM, in contrast to their predominant localization to the cytoplasmic leaflet of the parasite plasma membrane. In eukaryotic cells, phosphoinositides are predominantly localized to the cytoplasmic leaflet of cellular membranes, where they serve as spatially restricted membrane landmarks by recruiting and regulating cytosolic effector proteins through phosphoinositide-binding domains, including PH, PX, and FYVE domains [4, 40]. The detection of these phosphoinositides on both leaflets of the PVM therefore extends our understanding of PVM lipid organization beyond the phospholipids and glycolipids examined in our previous studies. In particular, these findings raise the possibility that the molecular interactions supported by phosphoinositides at the PVM differ from those at the parasite plasma membrane and other eukaryotic membranes. However, the present study does not establish whether phosphoinositides on the luminal leaflet interact with specific proteins or contribute directly to PVM function.

The molecular basis for the distribution of phosphoinositides across both leaflets of the PVM remains unclear. One possibility is that phospholipid scrambling during or shortly after parasite invasion allows lipids to redistribute between the two leaflets. Alternatively, continuous membrane remodeling during parasite replication or the activity of as-yet-unidentified lipid transport systems may contribute to the observed distribution. The involvement of lipid scramblases is an attractive hypothesis, but direct experimental evidence for their participation in phosphoinositide redistribution at the PVM is currently lacking. Further investigation will be required to identify the mechanisms responsible for the observed transbilayer distribution.

The observed distribution may have implications for the molecular organization and function of the PVM. Phosphoinositides recruit proteins involved in membrane trafficking, cytoskeletal organization, and signal transduction, thereby contributing to the organization of membrane-associated processes [4]. Their detection on both leaflets of the PVM raises questions about whether phosphoinositides on the luminal leaflet are accessible to parasite-derived proteins or other molecules within the vacuolar compartment, and whether their distribution influences the organization of proteins at the host–parasite interface. These possibilities are particularly relevant to understanding how the PVM establishes and maintains its specialized molecular environment. Nevertheless, the present study provides structural evidence of phosphoinositide distribution rather than direct evidence of its functional consequences. The potential roles of these lipids in nutrient transport, host immune evasion, and maintenance of the intracellular niche therefore remain to be investigated.

Importantly, the present findings extend our previous QF-FRL analyses of membrane lipid distribution, which demonstrated alterations in the transbilayer asymmetry of phosphatidylserine (PtdSer), phosphatidylethanolamine (PtdEtn), and the glycolipid GM3 in the PVM [41]. Together, these studies indicate that the unusual transbilayer organization of the PVM encompasses multiple classes of membrane lipids, including major phospholipids, a glycolipid, and phosphoinositides. The present study adds a new dimension to this picture by showing that the distribution of two signaling lipids, PtdIns(4)P and PtdIns(4,5)P_2_, also differs between the parasite plasma membrane and the PVM. This broader pattern suggests that PVM formation is associated with extensive remodeling of membrane lipid organization, although the mechanisms and functional consequences of these changes remain unresolved.

Whether similar phosphoinositide distributions occur in the PVMs of other apicomplexan parasites remains an important question. Comparative analyses of PVM lipid organization in *T. gondii* and *Plasmodium falciparum* may help determine whether the observed distribution is conserved among apicomplexans or reflects parasite-specific mechanisms. Such studies could further clarify how membrane lipid organization contributes to the formation and maintenance of the specialized host–parasite interface.

In conclusion, this study provides a leaflet-specific ultrastructural analysis of PtdIns(4)P and PtdIns(4,5)P_2_ in *T. gondii*. By demonstrating that both phosphoinositides are predominantly localized to the cytoplasmic leaflet of the parasite plasma membrane but are detected on both leaflets of the PVM, our findings extend previous observations of altered lipid asymmetry in the PVM to phosphoinositides. These results provide a structural basis for future investigations into the mechanisms governing PVM lipid organization and the potential consequences of this organization for phosphoinositide-dependent molecular interactions at the host–parasite interface.

## Data availability

All data generated or analyzed during this study are included in this published article.

## Supplemental Figure legends

**Fig. S1.**
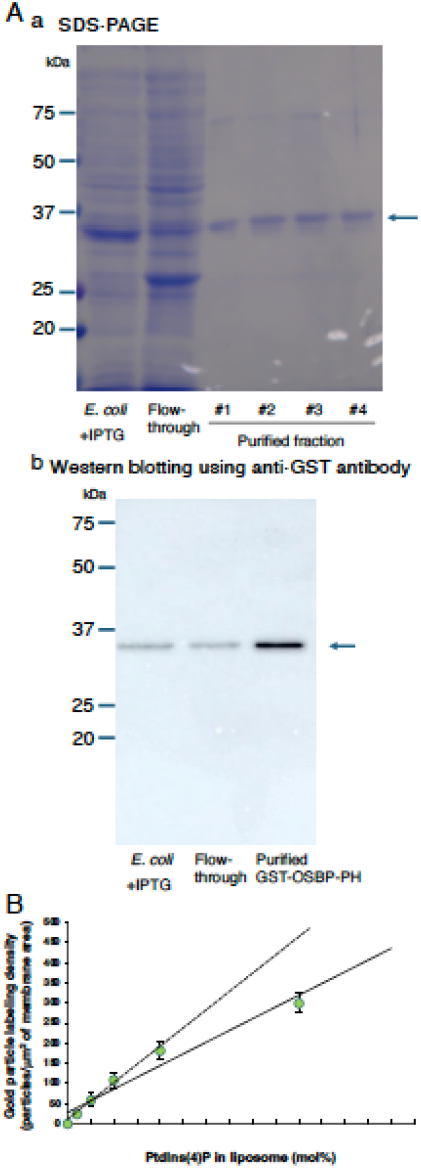
Validation of GST–OSBP-PH expression, purification, and PtdIns(4)P-dependent labeling. (A) SDS-PAGE (a) and Western blotting (b) of *E. coli* expressing GST–OSBP-PH after IPTG induction, the flow-through fraction collected after passage through a glutathione-Sepharose column, and four purified GST–OSBP-PH fractions (#1–#4). Western blotting was performed using an anti-GST antibody. Molecular mass standards are indicated on the left, and arrows on the right indicate the position of GST–OSBP-PH. (B) QF-FRL of unilamellar liposomes containing 0–5 mol % of PtdIns(4)P. GST-OSBP-PH was used at a concentration of 40 ng/mL. The number of colloidal gold particles per μm^2^ was measured in more than 90 randomly selected liposomes. The best-fitting lines were drawn by using 4 (0–2 mol %; dotted line) or 5 (0–5 mol %; solid line) points. The labeling density increased approximately linearly with the PtdIns(4)P content from 0 to 2 mol%, whereas a modest saturation tendency was observed at 5 mol%.

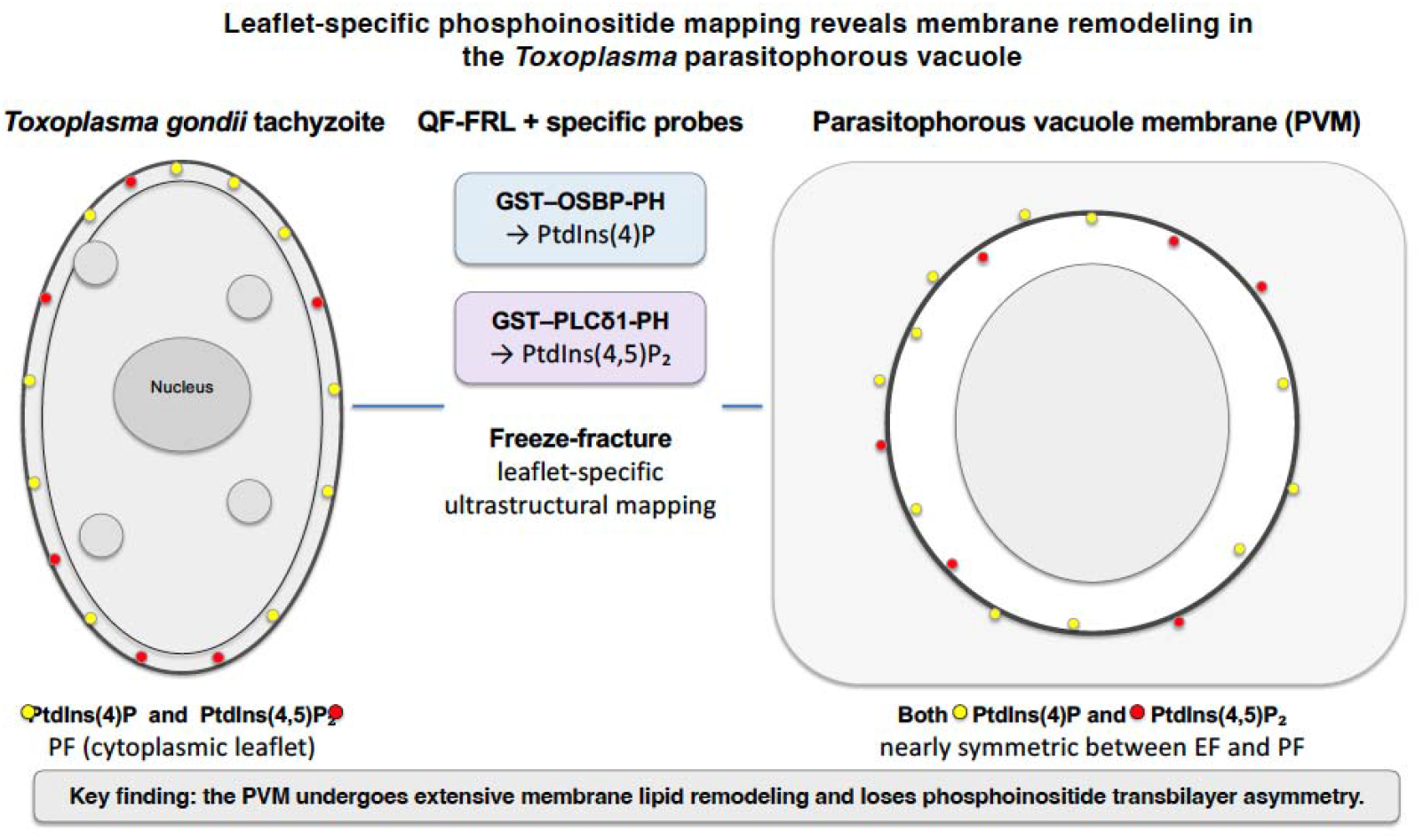

## References

[1] D.L. Daleke, Regulation of transbilayer plasma membrane phospholipid asymmetry, J Lipid Res 44(2) (2003) 233–42.

[2] A. Yamaji-Hasegawa, M. Tsujimoto, Asymmetric distribution of phospholipids in biomembranes, Biol Pharm Bull 29(8) (2006) 1547–53.

[3] G. van Meer, D.R. Voelker, G.W. Feigenson, Membrane lipids: where they are and how they behave, Nat Rev Mol Cell Biol 9(2) (2008) 112–24.

[4] T. Balla, Phosphoinositides: tiny lipids with giant impact on cell regulation, Physiol Rev 93(3) (2013) 1019–137.

[5] E.J. Dickson, B. Hille, Understanding phosphoinositides: rare, dynamic, and essential membrane phospholipids, Biochem J 476(1) (2019) 1–23.

[6] G.R. Hammond, M.P. Machner, T. Balla, A novel probe for phosphatidylinositol 4-phosphate reveals multiple pools beyond the Golgi, J Cell Biol 205(1) (2014) 113–26.

[7] G.R. Hammond, T. Balla, Polyphosphoinositide binding domains: Key to inositol lipid biology, Biochim Biophys Acta 1851(6) (2015) 746–58.

[8] L.D. Sibley, Invasion and intracellular survival by protozoan parasites, Immunol Rev 240(1) (2011) 72–91.

[9] E. Suss-Toby, J. Zimmerberg, G.E. Ward, Toxoplasma invasion: the parasitophorous vacuole is formed from host cell plasma membrane and pinches off via a fission pore, Proc Natl Acad Sci U S A 93(16) (1996) 8413–8.

[10] A.P. Sinai, Biogenesis of and activities at the Toxoplasma gondii parasitophorous vacuole membrane, Subcell Biochem 47 (2008) 155–64.

[11] A.M. Martin, T. Liu, B.C. Lynn, A.P. Sinai, The Toxoplasma gondii parasitophorous vacuole membrane: transactions across the border, J Eukaryot Microbiol 54(1) (2007) 25–8.

[12] B. Clough, E.M. Frickel, The Toxoplasma Parasitophorous Vacuole: An Evolving Host-Parasite Frontier, Trends Parasitol 33(6) (2017) 473–488.

[13] K.J. Saliba, K. Kirk, Nutrient acquisition by intracellular apicomplexan parasites: staying in for dinner, Int J Parasitol 31(12) (2001) 1321–30.

[14] A. Arabiotorre, M. Formanowicz, V.A. Bankaitis, A. Grabon, Phosphatidylinositol-4-phosphate signaling regulates dense granule biogenesis and exocytosis in Toxoplasma gondii, bioRxiv (2023).

[15] C.W. McNamara, M.C. Lee, C.S. Lim, S.H. Lim, J. Roland, O. Simon, B.K. Yeung, A.K. Chatterjee, S.L. McCormack, M.J. Manary, A.M. Zeeman, K.J. Dechering, T.S. Kumar, P.P. Henrich, K. Gagaring, M. Ibanez, N. Kato, K.L. Kuhen, C. Fischli, A. Nagle, M. Rottmann, D.M. Plouffe, B. Bursulaya, S. Meister, L. Rameh, J. Trappe, D. Haasen, M. Timmerman, R.W. Sauerwein, R. Suwanarusk, B. Russell, L. Renia, F. Nosten, D.C. Tully, C.H. Kocken, R.J. Glynne, C. Bodenreider, D.A. Fidock, T.T. Diagana, E.A. Winzeler, Targeting Plasmodium PI(4)K to eliminate malaria, Nature 504(7479) (2013) 248–253.

[16] L. Dembele, X. Ang, M. Chavchich, G.M.C. Bonamy, J.J. Selva, M.Y. Lim, C. Bodenreider, B.K.S. Yeung, F. Nosten, B.M. Russell, M.D. Edstein, J. Straimer, D.A. Fidock, T.T. Diagana, P. Bifani, The Plasmodium PI(4)K inhibitor KDU691 selectively inhibits dihydroartemisinin-pretreated Plasmodium falciparum ring-stage parasites, Sci Rep 7(1) (2017) 2325.

[17] W. Daher, J. Morlon-Guyot, L. Sheiner, G. Lentini, L. Berry, L. Tawk, J.F. Dubremetz, K. Wengelnik, B. Striepen, M. Lebrun, Lipid kinases are essential for apicoplast homeostasis in Toxoplasma gondii, Cell Microbiol 17(4) (2015) 559–78.

[18] A. Arabiotorre, V.A. Bankaitis, A. Grabon, Regulation of phosphoinositide metabolism in Apicomplexan parasites, Front Cell Dev Biol 11 (2023) 1163574.

[19] A. Arabiotorre, M. Formanowicz, V.A. Bankaitis, A. Grabon, Phosphatidylinositol-4-phosphate signaling regulates dense granule biogenesis and maturation in Toxoplasma gondii, PLoS Pathog 22(7) (2026) e1014451.

[20] S. Singh, M.M. Alam, I. Pal-Bhowmick, J.A. Brzostowski, C.E. Chitnis, Distinct external signals trigger sequential release of apical organelles during erythrocyte invasion by malaria parasites, PLoS Pathog 6(2) (2010) e1000746.

[21] A. Budu, C.R. Garcia, Generation of second messengers in Plasmodium, Microbes Infect 14(10) (2012) 787–95.

[22] V.B. Carruthers, S.N. Moreno, L.D. Sibley, Ethanol and acetaldehyde elevate intracellular [Ca2+] and stimulate microneme discharge in Toxoplasma gondii, Biochem J 342 ( Pt 2)(Pt 2) (1999) 379–86.

[23] J.L. Lovett, N. Marchesini, S.N. Moreno, L.D. Sibley, Toxoplasma gondii microneme secretion involves intracellular Ca(2+) release from inositol 1,4,5-triphosphate (IP(3))/ryanodine-sensitive stores, J Biol Chem 277(29) (2002) 25870–6.

[24] H.E. Bullen, Y. Jia, Y. Yamaryo-Botte, H. Bisio, O. Zhang, N.K. Jemelin, J.B. Marq, V. Carruthers, C.Y. Botte, D. Soldati-Favre, Phosphatidic Acid-Mediated Signaling Regulates Microneme Secretion in Toxoplasma, Cell Host Microbe 19(3) (2016) 349–60.

[25] K. Chintaluri, B.D. Goulden, C. Clemenza, G. Saffi, E. Miraglia, G.R.V. Hammond, R.J. Botelho, The PH domain from the Toxoplasma gondii PH-containing protein-1 (TgPH1) serves as an ectopic reporter of phosphatidylinositol 3-phosphate in mammalian cells, PLoS One 13(6) (2018) e0198454.

[26] T.E. Rusten, H. Stenmark, Analyzing phosphoinositides and their interacting proteins, Nat Methods 3(4) (2006) 251–8.

[27] O.V. Vieira, R.E. Harrison, C.C. Scott, H. Stenmark, D. Alexander, J. Liu, J. Gruenberg, A.D. Schreiber, S. Grinstein, Acquisition of Hrs, an essential component of phagosomal maturation, is impaired by mycobacteria, Mol Cell Biol 24(10) (2004) 4593–604.

[28] A. Fujita, J. Cheng, M. Hirakawa, K. Furukawa, S. Kusunoki, T. Fujimoto, Gangliosides GM1 and GM3 in the living cell membrane form clusters susceptible to cholesterol depletion and chilling, Mol Biol Cell 18(6) (2007) 2112–22.

[29] A. Fujita, J. Cheng, K. Tauchi-Sato, T. Takenawa, T. Fujimoto, A distinct pool of phosphatidylinositol 4,5-bisphosphate in caveolae revealed by a nanoscale labeling technique, Proc Natl Acad Sci U S A 106(23) (2009) 9256–61.

[30] A. Fujita, J. Cheng, T. Fujimoto, Quantitative electron microscopy for the nanoscale analysis of membrane lipid distribution, Nat Protoc 5(4) (2010) 661–9.

[31] Y. Kurokawa, A. Yoshida, E. Fujii, K. Tomioku, H. Hayashi, K. Tanabe, A. Fujita, Phosphatidylinositol 4-phosphate on Rab7-positive autophagosomes revealed by the freeze-fracture replica labeling, Traffic 20(1) (2019) 82–95.

[32] R. Konishi, Y. Nakashima, T. Masatani, M. Asada, H. Hassan, K. Fukuda, S. Kuriyama, Y. Nishikawa, O. Kaneko, V.B. Carruthers, A. Fujita, Parasitophorous vacuole membranes of Toxoplasma gondii and Plasmodium falciparum lack the lipid asymmetry characteristic of host cell plasma membranes, Biochim Biophys Acta Biomembr 1869(1) (2026) 184567.

[33] A. Yoshida, M. Shigekuni, K. Tanabe, A. Fujita, Nanoscale analysis reveals agonist-sensitive and heterogeneous pools of phosphatidylinositol 4-phosphate in the plasma membrane, Biochim Biophys Acta 1858(6) (2016) 1298–305.

[34] Y. Yamakuchi, Y. Kurokawa, R. Konishi, K. Fukuda, T. Masatani, A. Fujita, Selective increment of phosphatidylserine on the autophagic body membrane in the yeast vacuole, FEBS Lett 595(17) (2021) 2197–2207.

[35] T. Fujimoto, K. Fujimoto, Metal sandwich method to quick-freeze monolayer cultured cells for freeze-fracture, J Histochem Cytochem 45(4) (1997) 595–8.

[36] A. Fujita, T. Fujimoto, N. Ozato-Sakurai, H. Suzuki, A method for efficient observation of intracellular membranes of monolayer culture cells by quick-freeze and freeze-fracture electron microscopy, J Electron Microsc (Tokyo) 61(6) (2012) 441–6.

[37] M. Muramoto, N. Mineoka, K. Fukuda, S. Kuriyama, T. Masatani, A. Fujita, Coordinated regulation of phosphatidylinositol 4-phosphate and phosphatidylserine levels by Osh4p and Osh5p is an essential regulatory mechanism in autophagy, Biochim Biophys Acta Biomembr 1866(4) (2024) 184308.

[38] J. Schindelin, I. Arganda-Carreras, E. Frise, V. Kaynig, M. Longair, T. Pietzsch, S. Preibisch, C. Rueden, S. Saalfeld, B. Schmid, J.Y. Tinevez, D.J. White, V. Hartenstein, K. Eliceiri, P. Tomancak, A. Cardona, Fiji: an open-source platform for biological-image analysis, Nat Methods 9(7) (2012) 676–82.

[39] A. Burrell, V. Marugan-Hernandez, R. Wheeler, F. Moreira-Leite, D.J.P. Ferguson, F.M. Tomley, S. Vaughan, Cellular electron tomography of the apical complex in the apicomplexan parasite Eimeria tenella shows a highly organised gateway for regulated secretion, PLoS Pathog 18(7) (2022) e1010666.

[40] Y. Posor, W. Jang, V. Haucke, Phosphoinositides as membrane organizers, Nat Rev Mol Cell Biol 23(12) (2022) 797–816.

[41] R. Konishi, Y. Nakashima, T. Masatani, M. Asada, H. Hassan, K. Fukuda, S. Kuriyama, Y. Nishikawa, O. Kaneko, V.B. Carruthers, A. Fujita, Parasitophorous vacuole membranes of Toxoplasma gondii and Plasmodium falciparum lack the lipid asymmetry characteristic of host cell plasma membranes, Biochim Biophys Acta Biomembr (2026) 184567.

